# Selective labeling supports 5-thiooxazole post-translational modification in bufferins

**DOI:** 10.64898/2025.12.15.693703

**Authors:** Guy Lippens, Yanyan Li, Françoise Jacob-Dubuisson, Svetlana Dubiley

## Abstract

Multinuclear nonheme iron-dependent oxidases (MNIOs) constitute one of the largest families of enzymes involved in natural product biosynthesis. Distinct MNIO subfamilies utilize molecular oxygen to catalyze a wide variety of complex peptide rearrangements, including β-carbon excision and heterocyclization. Highly homologous MNIOs have been proposed to install either oxazolone-thioamides or 5-thiooxazoles as cysteine post-translational modifications in the closely related bufferin and EGKCG families of peptide chalkophores. These alternative structures prompted discussion of the subtle mechanistic features of MNIO enzymes that might determine reaction outcome. Here, we combine uniform 15N labeling with cysteine-specific carbonyl 13C labeling to unambiguously assign 5-thiooxazoles as the cysteine modifications in bufferins. Together with the recent identification of 5-thiooxazoles in three members of the sister EGKCG family and the re-assignment of the cysteine modification in oxazolin, these findings confirm that closely related MNIOs catalyze identical post-translational modifications.

---

RiPPs, or ribosomally synthesized and post-translationally modified peptides, constitute a vast and functionally diverse class of natural products found across all kingdoms of life [1]. RiPP tailoring enzymes show remarkable versatility in installing post-translational modifications (PTMs) on precursor peptides [2], generating unusual molecular structures that often challenge current analytical methods for accurate structure identifications [3–5]. Multinuclear non-heme iron-dependent oxidative enzymes (MNIOs) are one large family of enzymes that install post-translational modifications (PTMs) on ribosomally produced peptides [6,7].

RiPP chalkophores whose biosynthesis involves MNIOs represent a large superfamily of copper-binding RiPPs comprising distinct families [8]. Although the post-translational modifications in the methanobactin family were ultimately clarified after several revisions [9], the nature of MNIO installed modifications in other RiPP chalkophore families remains controversial. The first described representative of the EGKCG family [8], oxazolin from *Haemophilus influenzae* [10], reported cysteine residues converted into oxazolone thioamides (Figure 1A, **2**), a PTM identical to that identified in methanobactins [9,11]. However, an alternative structure for another EGKCG family member was later proposed: bulbicupramide from *Microbulbifer* sp. was described as containing 5-thiooxazole rings (Figure 1A, **3**), although the authors acknowledged the inherent difficulty in distinguishing closely related structures based solely on chemical shift values [12]. Similarly, Buf1 from *Caulobacter vibrioides*, a member of the bufferin family of RiPP chalkophores [8], was reported to have its two central cysteines modified to 5-thiooxazole rings whereas the two distal, unmodified cysteines form a disulfide bond [13].

The MNIO tailoring enzymes that modify cysteines in peptides from bufferin and EGKCG families are from the same subfamily and are only distantly related to the MNIO involved in methanobactin biosynthesis [8]. The distinct structural outcomes produced by highly homologous enzymes was therefore questioned [7] and prompted debate regarding the subtle mechanistic features that may dictate reaction specificity.

An elegant strategy combining free-thiol alkylation with HMBC-based connectivity analysis was recently applied to define the post-translational modifications in fontiphorin from *Fontimonas thermophila* and gonophorin from *Neisseria gonorrhoeae*, two EGCKG-family members [14]. The method relies on alkylation of the sulfur moiety and the always delicate elimination of alternative structures based on the presence or absence of HMBC peaks with unknown long-range couplings [15]. This approach led to the assignment of 5-thiooxazole modifications in these compounds, and equally concluded the presence of 5-thiooxazoles in the previously described oxazolin [14]. Nonetheless, we sought a more direct method, and applied it to confirm independently the structure of the PTMs in Buf1.

Our direct and unambiguous NMR experiment to assign the PTMs in Buf1 of *C. vibrioides* combines uniform ^15^N labeling and selective ^13^C labeling of cysteine carboxyl groups in the precursor peptide. Based on the proposed biosynthetic pathway [7,13] whereby conversion of cysteine into 5-thiooxazole should leave the original carbonyl group intact, it leaves a ^13^CO-^15^N peptide bond whereby the latter nitrogen is the amide of the residue downstream of the modified cysteine. Conversely, the formation of oxazolone thioamide should incorporate the former carbonyl group into the oxazolone heterocycle [16,17], leaving a ^12^C carbonyl atom in the peptide bond with the downstream residue (Figure 1A). Rather than detecting the ^13^C nucleus *via* long-range couplings to known proton signals in an HMBC experiment, we exploit the direct ^15^N-^13^CO ^1^J coupling to monitor the presence or absence of a ^13^C neighbor. The simplest experiment to make this distinction is the ^1^H, ^15^N plane of a HNCO experiment, commonly used in protein NMR of ^15^N,^13^C uniformly labeled samples. In the case of an unmodified cysteine residue (**1**) or of a 5-thiooxazole (**3**), one should see a peak in the HN(CO) plane for their downstream residue. In contrast, magnetization transfer between the carbonyl and ^15^N amide should be absent if the former is a ^12^C atom because the ^13^C carbonyl carbon is incorporated into the oxazolone ring, as is the case in structure (**2**). The value of the ^1^J coupling varies from -12 to -17Hz according to the residue type [18], but is not determined by any conformational factor [19] that could position it outside of this range.

To test this approach experimentally, we produced uniformly ^15^N-labeled Buf1, which was additionally selectively labeled with ^13^C at the carbonyl group of cysteines. We recorded the plane of an HNCO experiment [20], which corresponds to a ^1^H, ^15^N plane with filtering through the ^13^C carbonyl of the preceding residue: only ^1^H, ^15^N amide groups that are preceded by a ^13^C-labeled carbonyl can be detected in such an experiment. We therefore anticipated signals corresponding to the two unmodified cysteines (Cys12 and Cys52) visible through the amide correlations of their downstream residues, Tyr13 and Val53. For the modified Cys23 and Cys31, the presence or absence of cross-peaks for Lys24 and Ala32 would discriminate between the two possible PTMs.

The NMR plane of the HN(CO) experiment of the peptide displayed cross peaks for four, and not two, amide moieties (Figure 1B). Based on the previous assignment of the ^1^H, ^15^N spectrum [13], these four peaks correspond to the four residues that immediately follow cysteines in the primary sequence. The detection of signals for Lys24 and Ala32, together with those for Tyr13 and Val53, indicates that the carbonyl groups of Cys23 and Cys31 contain a ^13^C nucleus. These results clearly support the presence of two 5-thiooxazole PTMs in Buf1.

Combining the uniform ^15^N labeling and selective ^13^CO Cys labeling with the HN(CO) experiment to probe the presence or absence of a ^13^CO-^15^N peptide bond presents a direct method to evaluate the fate of the cysteine carbonyl carbon. Taken together with the recently published studies on other MNIO modified RiPPs [12,14], our results provide a definitive structural assignment for Buf1, and settle the apparent discrepancy in the products generated by highly homologous MNIO enzymes of the EGKCG and bufferin families of RiPP chalkophores.

**Figure 1.**
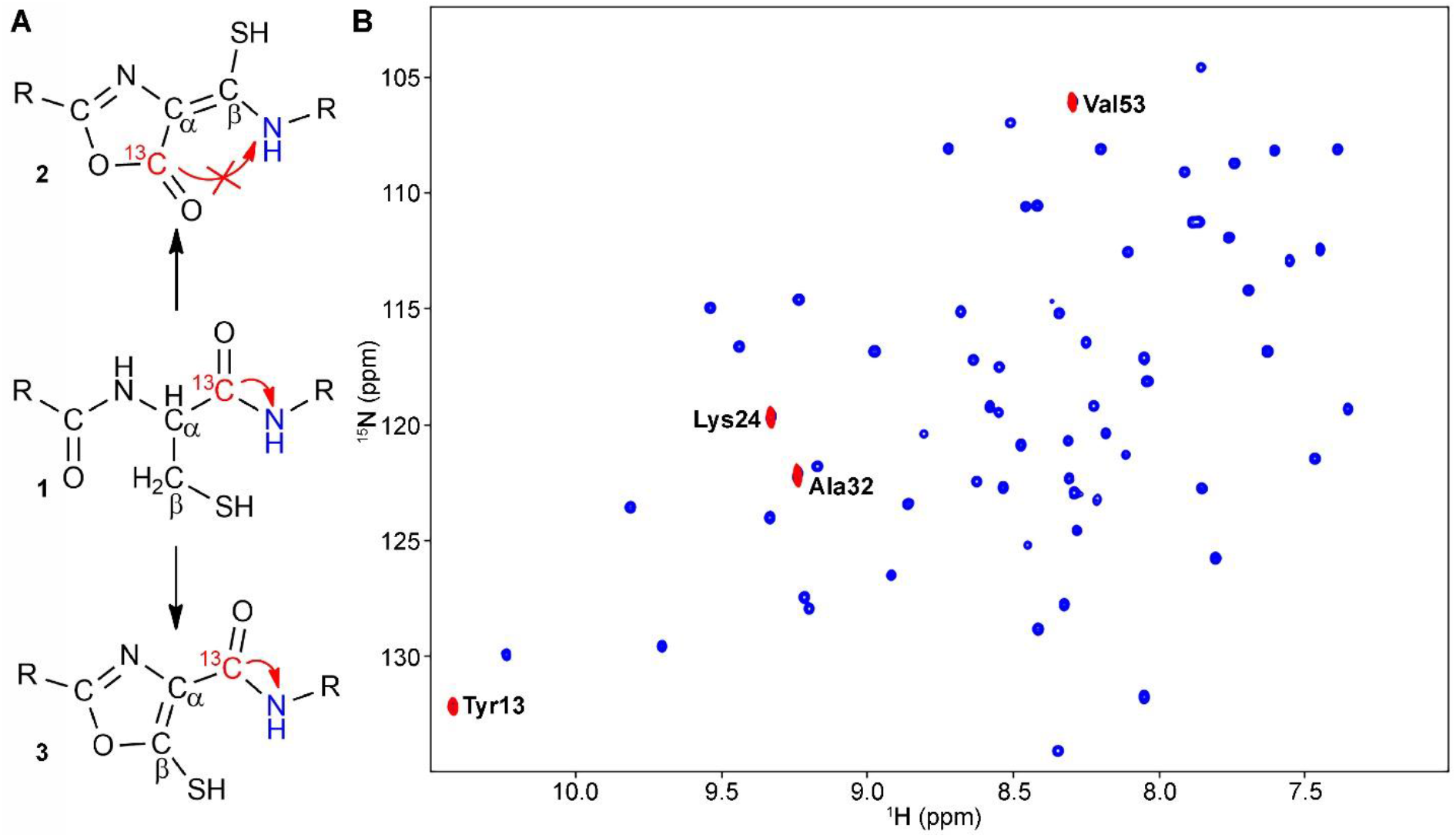
**A**. Schematic representation of possible MNIO-catalyzed posttranslational modifications of the cysteine residue (**1**), leading to a an oxazolone thioamide structure (**2**) with the ^13^C label in the oxazolone heterocycle, or a 5-thiooxazole ring (**3**), with the ^13^CO label connected by a ^1^J coupling constant to the ^15^N nucleus of the downstream residue. Only in structures **1** and **3** should the ^1^J ^15^N-^13^CO coupling give rise to a cross peak in the HN(CO) spectrum at the position of the downstream neighbor of the (modified) cysteine. **B**. ^1^H, ^15^N HSQC spectrum (blue) and HN(CO) plane (red) confirming that the ^13^C label is maintained at the carbonyl position of the modified Cys23 and Cys31.

## Methods

A cysteine-auxotrophic strain, *Escherichia coli* BW25113 ΔcysE [21], carrying the pCA24-buf1AstrBCD vector [13], was used to produce isotopically labeled Strep-tagged Buf1 from *C. vibrioides*. Cells were grown in M9 minimal medium supplemented with 0.5% glycerol, 1× MEM vitamin mixture, 1 g/L ^15^NH_4_Cl, and 0.25 mM cysteine-1-^13^C at 37 °C until the OD_600_ reached 0.6. Expression was then induced with 1 mM isopropyl β-D-thiogalactopyranoside, and cultures were incubated at 20 °C for 24 h. Harvested cells were resuspended in Buffer A (20 mM Tris, pH 7.5, 300 mM NaCl, 1 mM phenylmethylsulfonyl fluoride) and disrupted by sonication. The lysate was cleared by centrifugation, filtered through a 0.22 µm polyethersulfone membrane, and loaded onto a 1-mL StrepTactin Superflow column (IBA, Germany). After washing with 20 mL Buffer A, the protein was eluted with 2.5 mM desthiobiotin in Buffer A. Protein-containing fractions were pooled and buffer-exchanged into 50 mM sodium phosphate, pH 7.5 using Amicon centrifugal filters (3 kDa cutoff).

All NMR experiments were acquired at 293K on a 800 MHz spectrometer equipped with a 5-mm cryoprobe with pulsed field z-gradients. The sample contained ∼200µM of U-15N, Cys 13CO labeled bufferin1 in 50 mM sodium phosphate, pH 7.5. The data presented were recorded using the standard Bruker library pulse sequences: 2D 1H-15N HSQC and HNCO. All spectra were processed using Topspin 4.0.8 (Bruker Biospin). Spectral dimensions for the 2D 1H-15N HSQC and HN(CO) experiments were 15.22 × 38ppm, with sampling durations of 160 ms (t3) and 10.9 ms (t2). Spectra were centered at 4.7 ppm (1H) and 119 ppm (15N) setting the 13CO carrier at 173 ppm.

## Author contributions

All authors obtained, analyzed and interpreted the data, and wrote the manuscript collectively.

## Acknowledgments

This project was funded by the Agence Nationale de la Recherche (ANR) grants CuRiPP (ANR-22-CE44-0001-02) and MetalloRIPP (ANR-25-CE44-1111-01). S. D. was supported by a grant of the INRAE EXPLOR’AE program, a part of the France 2030 initiative, grant ANR-24-RRII-0003. We thank MetaboHub-MetaToul (Metabolomics and Fluxomics facilities, Toulouse, France), part of the French National Infrastructure for Metabolomics and Fluxomics, for access to the NMR facility.

## TOC graphic

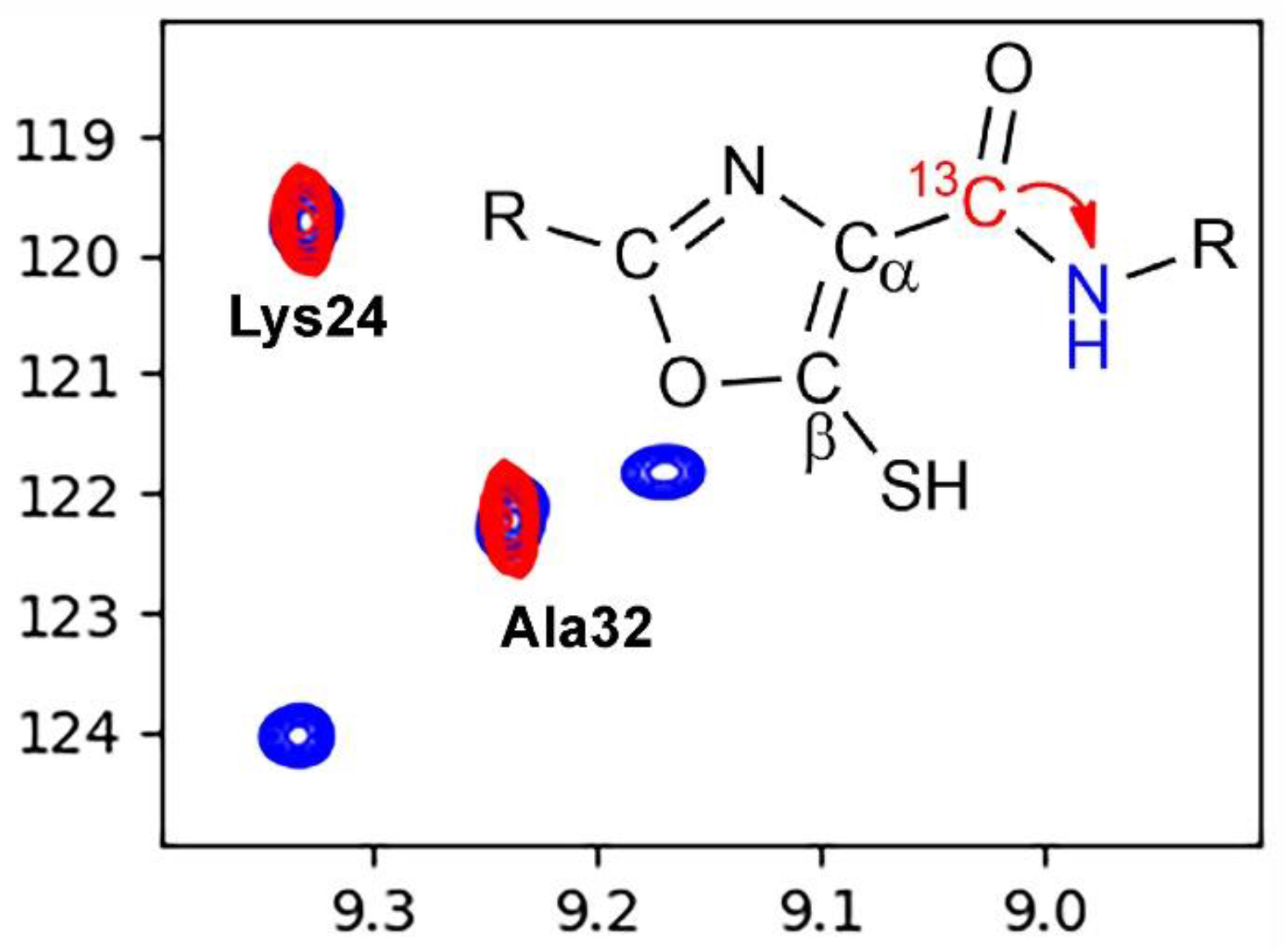

